# Resolving outbreak dynamics using Approximate Bayesian Computation for stochastic birth-death models

**DOI:** 10.1101/215533

**Authors:** Jarno Lintusaari, Paul Blomstedt, Tuomas Sivula, Michael U. Gutmann, Samuel Kaski, Jukka Corander

## Abstract

Earlier research has suggested that Approximate Bayesian Computation (ABC) makes it possible to fit simulator-based intractable birth-death models to investigate communicable disease outbreak dynamics with accuracy comparable to that of exact Bayesian methods. However, recent findings have indicated that key parameters such as the reproductive number *R* may remain poorly identifiable. Here we show that the identifiability issue can be resolved by taking into account disease-specific characteristics of the transmission process in closer detail. Using tuberculosis (TB) in the San Francisco Bay area as a case-study, we consider the situation where the genotype data are generated as a mixture of three stochastic processes, each with their distinct dynamics and clear epidemiological interpretation.

The ABC inference yields stable and accurate posterior inferences about outbreak dynamics from aggregated annual case data with genotype information. We also show that under the proposed model, the infectious population size can be reliably inferred from the data. The estimate is approximately two orders of magnitude smaller compared to assumptions made in the earlier ABC studies, and is much better aligned with epidemiological knowledge about active TB prevalence. Similarly, the reproductive number *R* related to the primary underlying transmission process is estimated to be nearly three-fold compared with the previous estimates, which has a substantial impact on the interpretation of the fitted outbreak model.

## 1 Introduction

Birth-death processes are flexible models used for numerous purposes, in particular for characterizing spread of infections under the so called Susceptible-Infectious-Removed (SIR) formulation of an epidemic process [Anderson and May, 1992]. Under circumstances where an outbreak of a disease occurs, but daily, weekly or even monthly incidence counts are not directly applicable or even available, the estimation of key epidemiological parameters, such as the reproductive number *R*, has to be based on alternative sources of information. This can be the case when the disease demonstrates large variability between the times of infection and onset, such as with *Mycobacterium tuberculosis*, or in retrospective analyses where all the information is no longer available. In such cases aggregate measures of the clusteredness of cases, for instance by genotype fingerprints, can be used as alternative source of information. However, the likelihood-based inference is often considerably more challenging in these cases compared to standard outbreak investigations relying solely on incident count data.

As a solution to such a setting, Tanaka et al. [2006] proposed fitting birth-death (BD) models to tuberculosis (TB) outbreak data using Approximate Bayesian Computation (ABC). Later on the same setting was used in numerous ABC studies while the ABC methodology was being developed [see e.g. Sisson et al., 2007, Blum, 2010, Fearnhead and Prangle, 2012, Del Moral et al., 2012, Baragatti et al., 2013, Albert et al., 2015]. Stadler [2011] and Aandahl et al. [2014] also tested the ABC procedure against an exact Bayesian inference method based on elaborate Markov Chain Monte Carlo (MCMC) sampling scheme. These investigations considered TB outbreak data from San Francisco Bay area originally collected by Small et al. [1994], who reported results from extensive epidemiological linking of the cases, as well as the corresponding classical IS6110 fingerprinting genotypes. Such genetic data from the causative agent *Mycobacterium tuberculosis* are natural to characterize using the infinite alleles model (IAM), where each mutation is assumed to result in a novel allele in the bacterial strain colonizing the host. When lacking precise temporal information about the infection and the onset of the active disease, the numbers and sizes of genotype clusters can be used to infer the parameters of the BD model as shown by Tanaka et al. [2006], Aandahl et al. [2014].

Lintusaari et al. [2016] raised an issue of non-identifiability of *R* for the TB outbreak model in cases when both the birth and death rates were unknown in the underlying birth-death process. This was demonstrated as a nearly flat approximate likelihood over the parameter space of *R*. Also it was found that in cases when *R* was identifiable, the acquired estimate was dependent on the assumed population size *n*, and correspondingly, by fixing *R* it was possible to infer the infectious population size n. In the earlier investigations by Tanaka et al. [2006] it was concluded that a large infectious population size *n* = 10000 was required for the BD simulator to produce similar levels of genetic diversity as observed in the San Francisco Bay data. However, so large a population of active TB disease cases is in stark contrast with the existing epidemiological evidence [Small et al., 1994].

Here we introduce an alternative formulation of the BD model which resolves the identifiability issue of *R*, and allows simultaneously for the estimation of the underlying infectious population size *n*. Our model incorporates epidemiological knowledge about the TB infection and disease activation processes by assuming that the observed genotype data represent a mixture of two birth-death processes, each with clearly distinct characteristics. By evaluating the ABC inference results of our model in the light of the epidemiological information available from Small et al. [1994], it is seen that both the significantly reduced infectious population size *n* and the increased *R* for the main driver component of the model make good sense and drastically change the interpretation of the fitted model in comparison to the earlier ABC studies.

In the new model we consider latent and active TB infections separately, as only the latter may lead to new transmission events. Transmission clusters are formed by a recent infection that rapidly progresses to an active TB and is spread further in the host population. Due to the rapid onset, the fingerprint of the pathogen remains the same in the transmission process and the patients consequently form an epidemiological cluster. If, on the other hand, the infection remains latent, the pathogen will undergo mutations and hence alters its fingerprint over the years [Small et al., 1994]. By this and other epidemiologically motivated modelling choices we show that the model becomes identifiable. Due to the rather modest requirements for the available data and flexibility of modelling in ABC, our BD model can be applied to many similar settings beyond the case study considered in this article.

## 2 The BD model for TB epidemic

The new model is based on the birth-death (BD) process where birth events correspond to an appearance of a new case with an active TB. A death event corresponds to any event that makes the existing host non-infectious, such as death, sufficient treatment, quarantine or relocation away from the community under investigation. The model incorporates two BD processes and one pure birth process that have a epidemiologically based interpretation. As in the standard BD process, the events are assumed to be independent of each other and to occur at specific rates. The time between two events is assumed to follow the exponential distribution specified by the rate of occurrence, causing the number of events to follow the Poisson distribution. The time scale considered here is one calendar year. The evolution of an infectious population is simulated by drawing events according to their rates.

Building upon the BD process, the simulated population carries auxiliary information. At birth, a case is assigned a cluster index that represents the specific genetic fingerprint of the pathogen and detemines the cluster the case belongs to. The simulated output includes the cluster indexes that are recorded when cases become observed. Next we explain the model in more detail and notify differences to the model of Tanaka et al. [2006].

First, we assume that observations are collected within a given time interval that matches the observed data. In the case of the San Francisco Bay data, the length of this interval is two years [Small et al., 1994]. The observations are collected from the simulated process after a sufficient warm-up period, so that the process can be expected to have reached stable properties (exemplified in Figure 1). A patient becomes observed with probability *p_obs_* when they cease to be infectious, i.e. when they undergo a death event in the simulation. Combining both being observed and ceasing to be infectious under the death event is based on the assumption that a typical patient is treated promptly after being diagnosed [Sreeramareddy et al., 2009]. In contrast to the model of Tanaka et al. [2006], there is then no separate observation sampling phase nor a prior estimate for the underlying population size.

**Figure 1:**
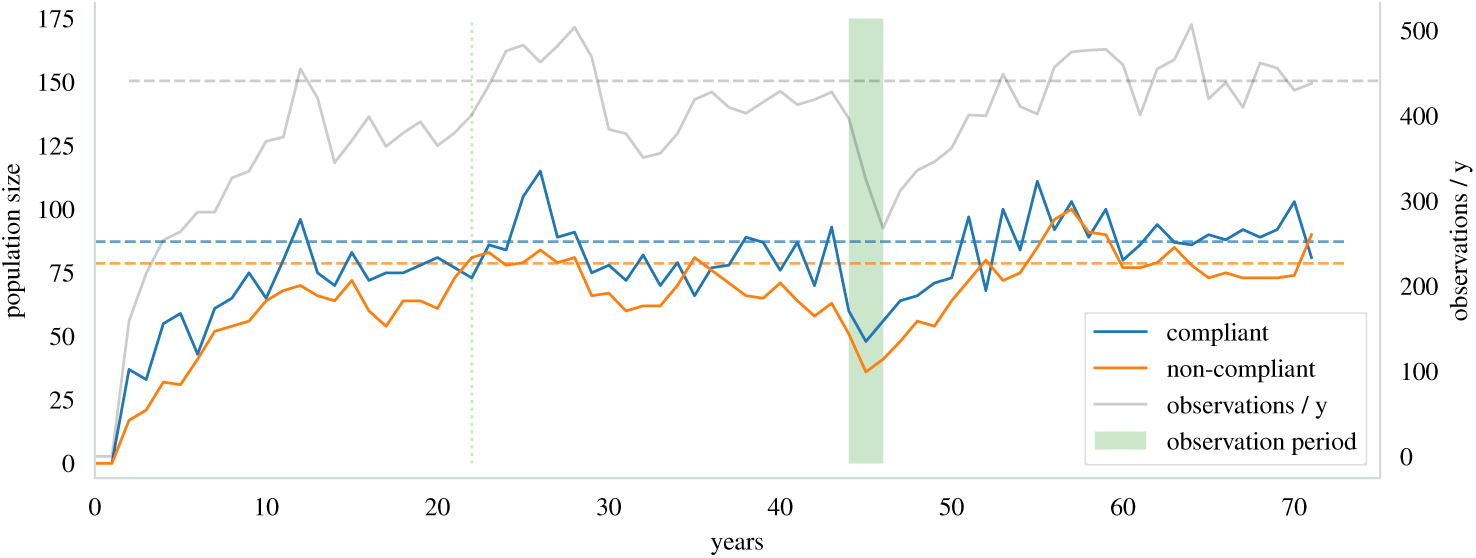
An illustration of simulated compliant and non-compliant populations as observed in the end of each year. The dashed lines are the balance values. The population sizes fluctuate around them after the process has matured. Both populations have surpassed their balance value at least once after 22 years. The observation period is the green patch. The grey line shows the number of observations that would have been collected within each year in the simulation. The number of observations from the observation period together with the clustering structure of the observations are used in the inference of the epidemiological parameters.

Second, a burden rate parameter *β* is introduced to reflect the rate at which new active TB cases with an previously unseen fingerprint of the pathogen appear in the community. This is the pure birth process of the model and reflects the reactivations of TB from the underlying latently infected population and immigration. In the simulation such cases receive a new cluster index that has not been assigned to any earlier case. Unlike Tanaka et al. [2006], mutations are not explicitly modelled, but are assumed to occur during the latent phase of infection over the years [Small et al., 1994].

Third, two distinct birth-death processes are introduced for cases that are either *compliant* or *non-compliant* with treatment. The birth-death processes are parametrized with birth rates *τ_i_*, and death rates *δ_i_*, where subscript *i* = 1 denotes the non-compliant population and *i* = 2 the compliant population. As noted in Small et al. [1994], a significant factor behind the largest clusters in the observed data were non-compliant patients who stayed infectious for several months and belonged to subgroups under increased risk of rapid development of active TB due to conditions such as AIDS or substance abuse. Typical patients who are compliant with the therapy cease to be infectious relatively fast and do not transmit the disease as effectively before their diagnosis and treatment. Meta-analysis of typical time delays before diagnosis can be found from Sreeramareddy et al. [2009].

We assume that a new TB case is non-compliant with therapy with probability *p*_1_. At transmission (birth event) in the simulation this probability is used to determine the population the new case belongs to. We also assume that the epidemic is in a steady state (Figure 1) by requiring that compliant cases have a reproductive number *R*_2_ = *τ*_2_/*δ*_2_ < 1 below one and that the reproductive number *R*_1_ of the non-compliant cases is constrained such that the population does not grow without limit. We will next identify the subspace of parameter values that conform to this assumption.

## 3 Statistical Analysis

Let subscript *i* = 1 denote a parameter of the non-compliant population and *i* = 2 the compliant population. The sizes of the compliant and non-compliant subpopulations can be analyzed by investigating the parameters of the three birth-death processes in the proposed model. First we notice that a size of a subpopulation follows a compound birth-death process whose birth-rate is a linear function of the burden rate and the birth rates of the two patient type populations at their respective sizes. For instance the birthrate of the non-compliant subpopulation is *p*_1_(*β* + *τ*_1_*n*_1_ + *τ*_2_*n*_2_) where *n*_1_ and *n*_2_ are the current sizes of the subpopulations and *p*_1_ is the probability of a case being non-compliant. The corresponding death rate is *δ*_1_*n*_1_. Using this approach we can determine the balance sizes *b*_1_ and *b*_2_ of the subpopulations, meaning values of *n*_1_ and *n*_2_ for which the birth and death rates of both of the subpopulations are equal. This corresponds to a state where the subpopulation sizes neither shrink or grow. The balance values *b*_2_ and *b*_1_ are obtained by solving the following set of linear equations:

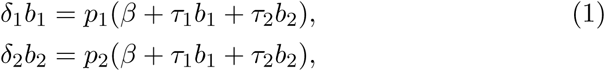
where *p*_2_ = (1 − *p*_1_) is the probability of a new case being compliant. The linear equations yield the following solution

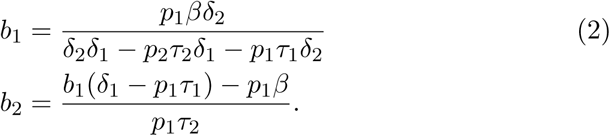

Given this solution, the balance values *b*_1_ and *b*_2_ exist when

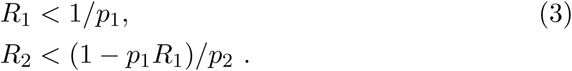

Assuming for instance that *p*_2_ = .95 [Small et al., 1994], this translates to *R*_1_ < 20.

Equation 2 allows also one to approximate the mean number of observed cases per year by defining the approximation as

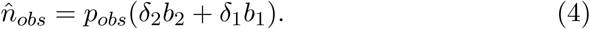

Figure 1 illustrates how the population sizes fluctuate near their balance values in the simulation after a sufficient warm-up period.

### 3.1 Parameter inference

Approximate Bayesian computation was used to carry out the parameter inference due to the unavailability of the likelihood function. This is the same approach as used by Tanaka et al. [2006] with the original model. The result will be a sample from the approximate posterior distribution 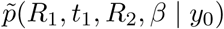 [see e.g. Lintusaari et al., 2017].

We used the Engine for Likelihood-Free Inference (ELFI) software [Lintusaari et al., titb] to perform the inference. We sampled 1000 parameter values with rejection sampling from a total of 6M simulations. A visualization of the ELFI model can be found from Figure S1 in Supplementary Material. The observed data are available in the article of Small et al. [1994]. Furthermore we have released the source code of the simulator and the corresponding ELFI model in GitHub^1^ that allow a replication of this study.

#### 3.1.1 Priors

We set priors over the burden rate *β*, reproductive numbers *R*_1_ and *R*_2_, and the net transmission rate *t*_1_ = *τ*_1_ − *δ*_1_ of the non-compliant population. For the compliant population the death rate is fixed to an estimate *δ*_2_ = 5.95 [Sreeramareddy et al., 2009, the total delay estimate] that can be transformed to a net transmission rate via *t*_2_ = *δ*_2_ (*R*_2_ − 1). Based on the details in Small et al. [1994] describing the San Francisco Bay data, the probability of becoming observed was fixed to *p_obs_* = 0.8 and the probability of a new case being non-compliant was set to *p*_1_ = 0.05.

The burden rate *β* is given an informative prior that is able to produce a sufficient number of clusters with respect to the observed data. Specifically, we set

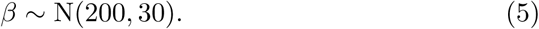

The net transmission rate *t*_1_ is given a uniform prior over a large interval from 0 to 30. Given the solution in Equation 3, the reproductive numbers *R*_1_ and *R*_2_ are given uniform prior over subspace that ensures the process has a steady state. More specifically

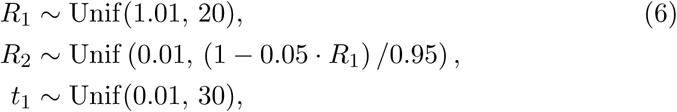

To optimize the computation given the observed data, we set the following additional constraints:

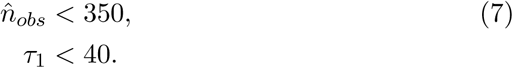

These constraints were checked to have a negligible effect on the acquired estimates. They however prevented simulations with extremely unlikely parameter values thus saving considerable amount of computation time. Effectively due to the constraints, the values of *R*_1_ were smaller than 15. Figure S2 in Supplementary Material shows samples drawn from the priors under these constraints.

#### 3.1.2 Summary statistics

The summary statistics used in earlier approaches [see e.g. Tanaka et al., 2006, Lintusaari et al., 2016] are not directly applicable to the proposed model. This is due to the differences between the models that cause for example the number of observations in the sample to vary rather than being fixed. However, the earlier summaries still provide a good starting point for developing a more comprehensive set of summaries.

We found the following eight summary statistics to be informative about the parameters. The first summary was the number of observations. Five of the summaries were related to the clustering structure: the total number of clusters, the relative number of singleton clusters, the relative number of clusters of size two, the size of the largest cluster and the mean of the successive difference in size among the four largest clusters (Table S1 in Supplementary Material).

In addition we included two summaries from observation times of the largest cluster. Observation times were not included in the earlier approaches and proved to be useful in identifying the net transmission rate *t*_1_. These were the number of months from the first observation to the last and the number of months when at least one observation was made. This data could be extracted from Figure 2 in Small et al. [1994].

**Figure 2:**
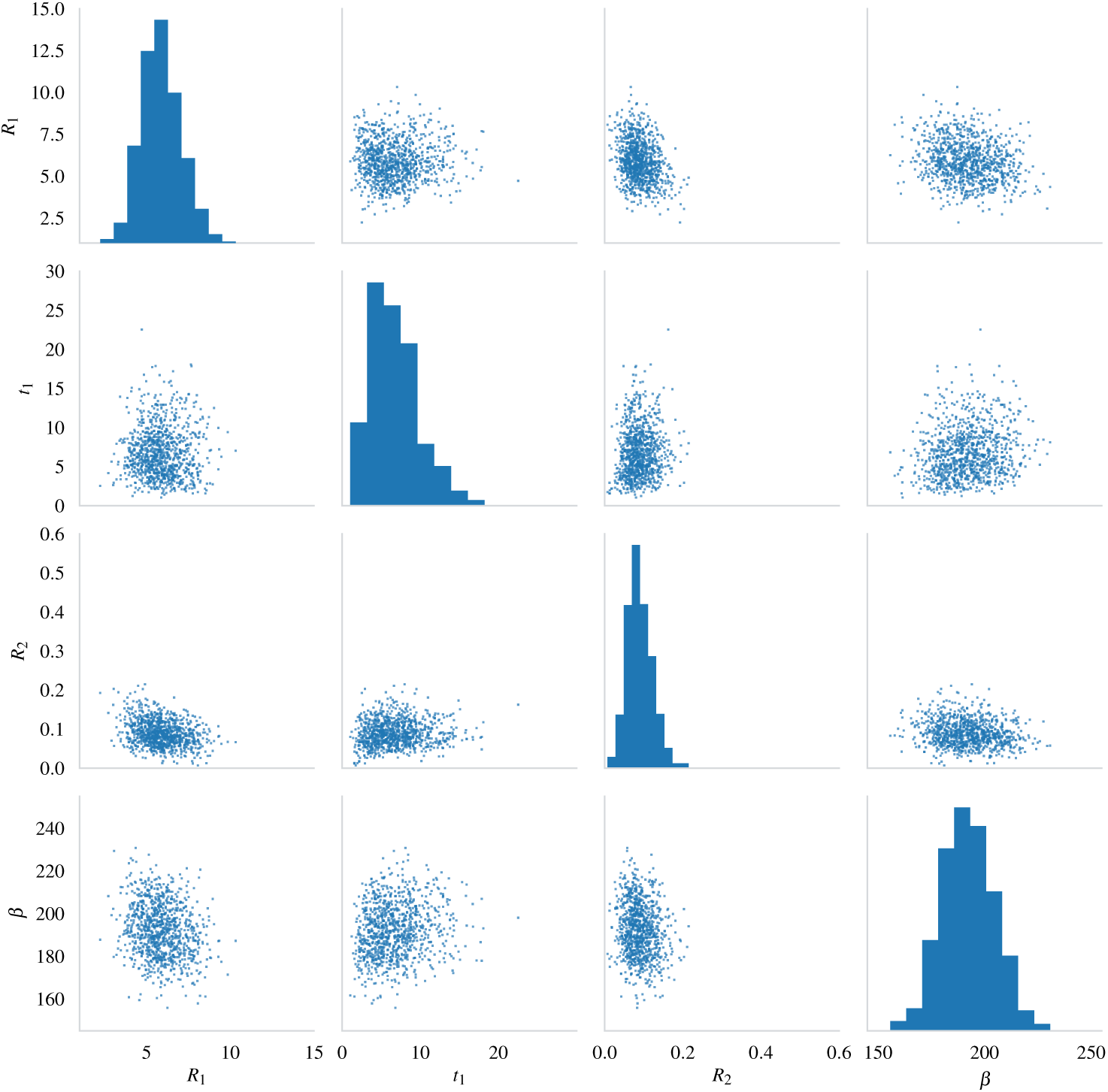
Posterior sample of size 1000 from the approximate posterior distribution 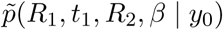 plotted as a scatter matrix. Compare to the prior in Figure S2 in Supplementary Material.

The distance function was the Euclidean distance between the weighted summary statistics of the observed and simulated data (Table S1 in Supplementary Material). The weights were chosen to adjust and even up differences in the magnitudes of the different summaries. The inference is not very sensitive to the exact values of these weights. The chosen values were found to perform well with respect to the evaluation of the model. The resulting threshold for the acquired sample was *ϵ* = 31.7 with the smallest distance being 12.5.

## 4 Results

Figure 2 shows a sample of 1000 values from the joint approximate posterior distribution 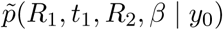. The pairwise sample clouds seem reasonably concentrated and are away from the edges of the axes and inside the support of the prior (compare to the prior in Figure S2 in Supplementary Material). The histograms and scatter plots look rather normally shaped, the only minor exception being the net transmission rate of the non-compliant population *t*_1_, that has a slight tail towards high values. The posterior suggests that the model is identifiable for the San Francisco dataset.

The posterior means, medians and 95% credible intervals are given in Table 1. The means and medians are close to each other supporting the above observation about normality. The *t*_1_ has the largest discrepancy due to its small tail mentioned above.

**Table 1:** Posterior summaries.

| Parameter | Mean | Median | 95% CI |
| --- | --- | --- | --- |
| $R_1$ | 5.88 | 5.79 | (3.68, 8.16) |
| $t_1$ | 6.74 | 6.25 | (1.57, 12.9) |
| $R_2$ | 0.09 | 0.09 | (0.03, 0.15) |
| $\beta$ | 192 | 192 | (170, 216) |

### 4.1 Evaluating the model identifiability

To further evaluate the reliability of the acquired estimates, we compute the mean and median absolute errors (MAE and MdAE) of the mean, and the coverage property (Wegmann et al, 2009), with 1000 synthetic observations from the posterior with known parameter values. These results include the ABC approximation error [see e.g Lintusaari et al., 2017] caused by the summary statistics and the threshold of 31.7.

Table 2 lists the MAE and MdAE with the 95% error upper percentile. These are useful in quantifying how much the estimate deviates from the actual value on average. The burden rate *β* and the reproductive number of the non-compliant population *R*_1_ have the smallest relative MAEs, 4.0% and 14.9%, respectively. The reproductive number *R*_2_ of the compliant population and the net transmission rate *t*_1_ of the non-compliant population have MAEs of 29.5% and 44.2%. The MAE of the latter seems rather high. Also the 95% percentile (Table 2) indicates that in 5% of the trials the error was substantial. Investigating the issue further showed that for some of the synthetic datasets, the net transmission rate parameter *t*_1_ was not identifiable, meaning that the synthetic data in those cases was not informative enough to produce a clear mode for the parameter. Also *R*_2_ suffered slightly from the same issue. This kind of situation where some of the synthetic datasets turn out uninformative is a rather common occurrence in cases where there is little data available. Because of these exceptions, MdAE might be a more appropriate measure as it is not as much influenced by the results of the non identifiable datasets in the trials. Relative MdAE errors were 21.9% and 32.1% respectively. Figure S3 in Supplementary Material visualizes the estimated values against their actual values for each of the parameters.

**Table 2:** Mean and Median Absolute Errors in 1000 trials with synthetic data from the posterior

| Parameter | MAE | Relative MAE<sup>2</sup> | MdAE | Relative MdAE | 95% percentile |
| --- | --- | --- | --- | --- | --- |
| $R_1$ | 0.85 | 14.9% | 0.72 | 12.6% | 2.00 |
| $t_1$ | 2.68 | 44.2% | 1.98 | 32.1% | 7.66 |
| $R_2$ | 0.024 | 29.5% | 0.018 | 21.9% | 0.07 |
| $\beta$ | 7.6 | 4.0 % | 6.1 | 3.1% | 19.8 |

The coverage property [Wegmann et al., 2009] is used to assess the reliability of the inference by checking whether the spreads of the acquired posterior distributions are accurate. Given a critical level *α*, the true parameter value should be outside the (1 − *α*) credible interval of the posterior with probability *α*. The estimated *α*-values from 1000 marginal posteriors with known true parameter values were satisfactory (Figure 3). For the critical level *α* = .05 the estimated a-values were 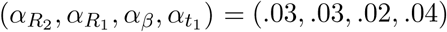. The overall performance with different *α* was similar to this case in the sense that *α_β_* suffered from a larger error compared to the estimates for the other parameters (Figure 3).

**Figure 3:**
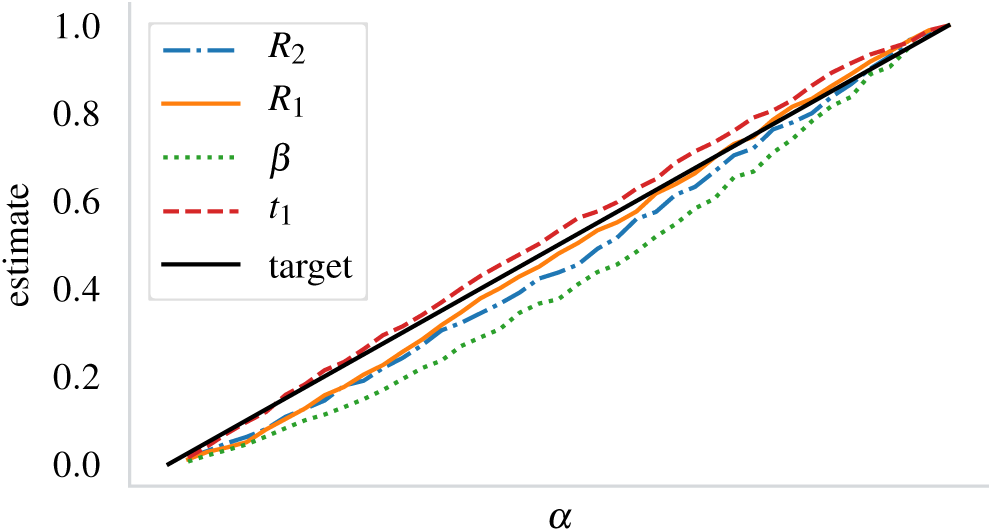
Estimates for the critical level *α* at different levels. The estimates are computed from 1000 synthetic datasets from the posterior. For the reference, the estimates for *α* = .05 were (.030, .028, .020, .041) in the same order as in the legend.

## 5 Discussion

We have proposed a stochastic birth-death model extending from several previous articles examining the use of simulator-based inference for the spread of active TB within a community. Outbreaks of TB are characterized by epidemiologically linked clusters of patients with active TB that emerge within a relative short time interval. The construction of the extended model was motivated by several epidemiological observations made by Small et al. [1994] concerning the San Francisco Bay transmission cluster data. Each of the largest clusters were largely formed by a non-compliant patient. In the largest cluster such a patient apparently infected 29 additional patients. The earlier approach [Tanaka et al., 2006, Aandahl et al., 2014] suffered from inability to reproduce these large clusters with an appropriate level of heterogeneity in the cluster sizes without a prior assumption of a very large underlying infectious population size (in the order of 10000) [Tanaka et al., 2006, Lintusaari et al., 2016]. Based on epidemiological knowledge about TB such a large infectious population size is extremely unlikely to have existed in the study region during the observation time interval. Furthermore it was shown that this assumption had a considerable effect on the estimate of the reproductive number *R*.

Under our new model, a prior estimate of the population size is not needed. Instead, the proposed model yields population size estimates as a by-product of the inference. For the San Francisco Bay data, the mean and median sizes were 48.4 and 48 for the compliant population and 13.5 and 11 for the non-compliant population.

The ability of the proposed model to estimate the population size parameters together with the other parameters follows from several important changes in the proposed model. One of them is that observations are collected during the observation period that matches the length of the actual observation period. In the original model observations were collected as a snapshot at a single point of time which essentially required that large clusters had to exist at that time to match the observed counts in the data. However, the observed counts are in reality a result of observations made over time as the outbreak evolves and no large clusters necessarily existed at any single point of time. Figure 2 in Small et al. [1994] shows how the patients were diagnosed at different times over the observation period. Furthermore the inclusion of non-compliant patient type in the model both more closely represents the description of Small et al. [1994] and naturally enables the formation of heterogeneity in the cluster sizes.

In the proposed model, the reproductive numbers represent the average number of infections that rapidly progress to active TB, caused by a single already infectious case. This counting therefore excludes infections remaining latent, which are instead indirectly captured via the burden rate parameter *β*. We estimate that the reproductive number of the non-compliant patients is *R*_1_ = 5.88 with the 95% credible interval (CI) (3.68,8.16) (Table 1). The estimate is nearly three-fold compared to the estimate of 2.10 in Aandahl et al. [2014] with the same data, which provided a single estimate for the whole infectious population without considering differences between patient types. The reproductive number of compliant cases is estimated to be *R*_2_ = 0.09 with a 95% CI (0.03, 0.15).

It should be noted here that being compliant or non-compliant are thought to characterize the type of a patient and the model decides this at the time of the birth event. In reality, the non-compliant cases are usually diagnosed (i.e. observed) earlier compared to when they cease to be infectious, which implies that the simulator model deviates slightly from typical observation processes in this respect. However, considering that this discrepancy applies to only 5% of all the observed cases, we do not expect any sizeable bias to arise from this assumption. Furthermore, the summary statistics used do not consider exact diagnosis times but rather just the span and the rate at which they occur.

The model identifiability was found to be satisfactory for the San Francisco Bay dataset (Figure 2). The average error in the estimate of *R*_1_ with the proposed method is evaluated to be 14.9% (0.85 in absolute terms, Table 2). The same for *R*_2_ is 29.5% (0.024 absolute), although the median error (21.9%, 0.018 absolute) is probably a more reasonable value due to the reasons discussed earlier. The coverage property analysis [Wegmann et al., 2009] suggests that the credible intervals provided by the model are sensible.

As the IS6110 typing remains in epidemiological use despite of advances in whole-genome sequencing of TB isolates, our model could be used for investigations in particular in middle and low income countries, where the TB burden is often also highest. For example, the estimates for the epidemiological parameters could be used to gain insight to the relative efficacy of the control programs across multiple communities. Given the apparent success by which the non-identifiability issue for *R* and the assumption of *a priori* known infectious population size were resolved by extending the BD model by relevant and often available epidemiological knowledge, it would be interesting to generalize the approach in the future to other pathogens for which the sampling process or other factors render the simulator-based inference as the most promising estimation method.

## Acknowledgements

We would like to acknowledge support for this project from the Academy of Finland (Finnish Centre of Excellence in Computational Inference Research COIN) and grants 294238, 292334 and the ERC grant no. 742158. We acknowledge the computational resources provided by the Aalto Science-IT project.

## Supplementary material

**Table S1:**
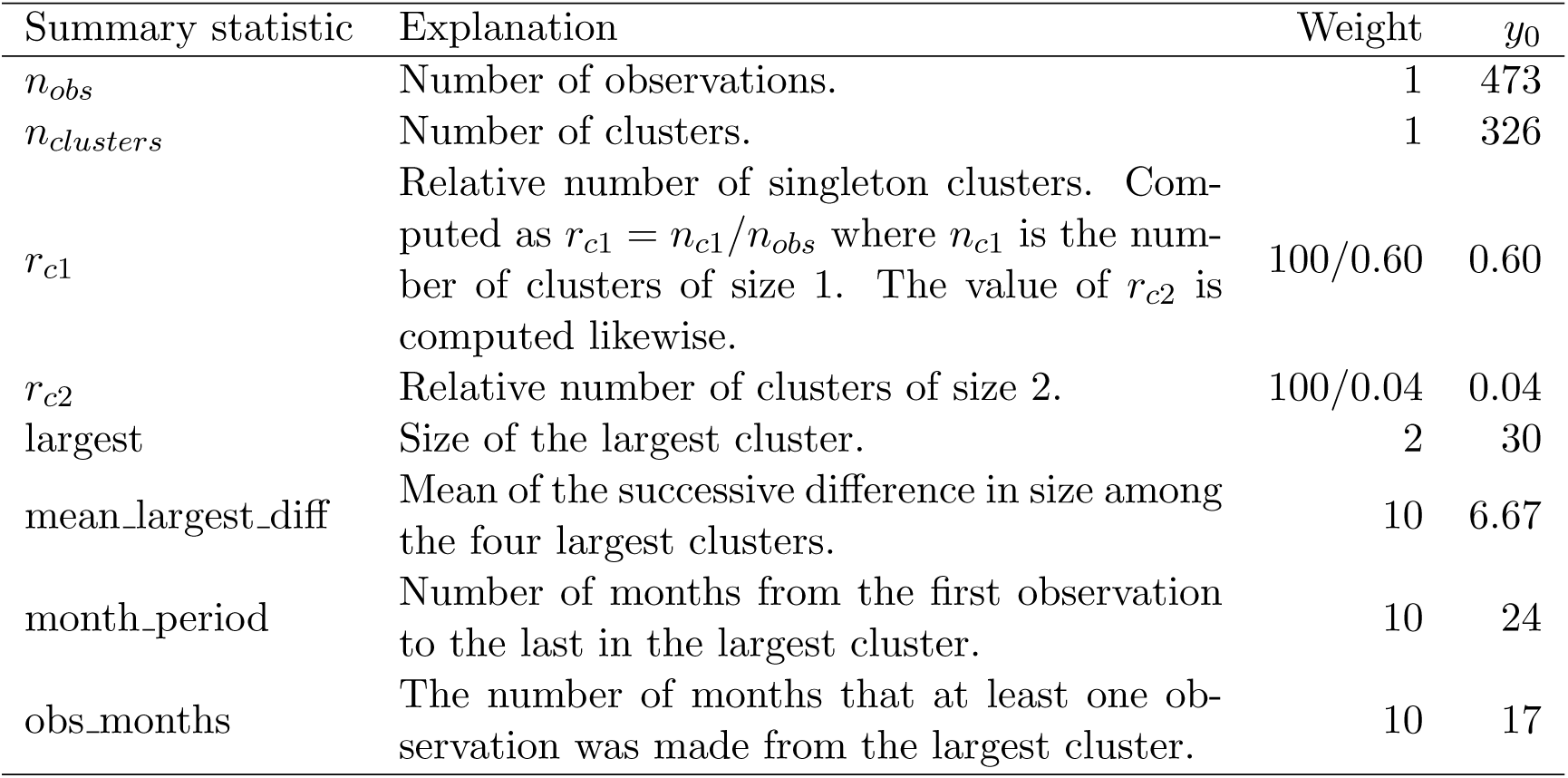
The summary statistics, their weights, and the values of the summary statistics for the observed data *y*_0_.

**Figure S1:**
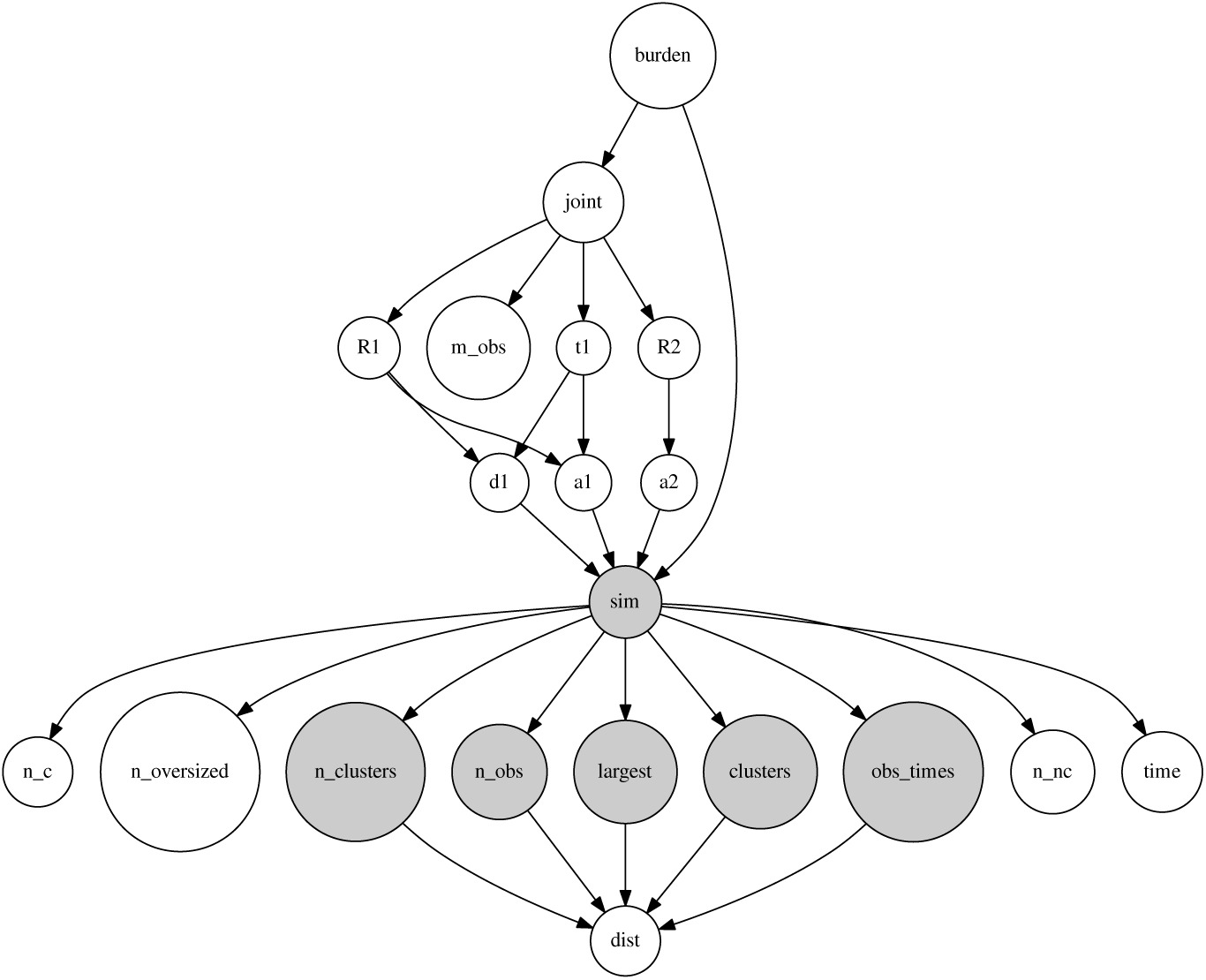
The ELFI model used in computing the posterior as visualized by ELFI. The grey node named *sim* is the simulator. It takes in the rate parameters that are transformed from the parameters of interest. The grey child nodes are summary statistics or quantities from which the summary statics are computed from (not all of the summaries in Table S1 were available as individual nodes but were computed in *dist* from *obs_times* and *clusters*). Some other side information was also collected, such as the total simulated time and size of the compliant and non-compliant populations at the end of the simulation.

**Figure S2:**
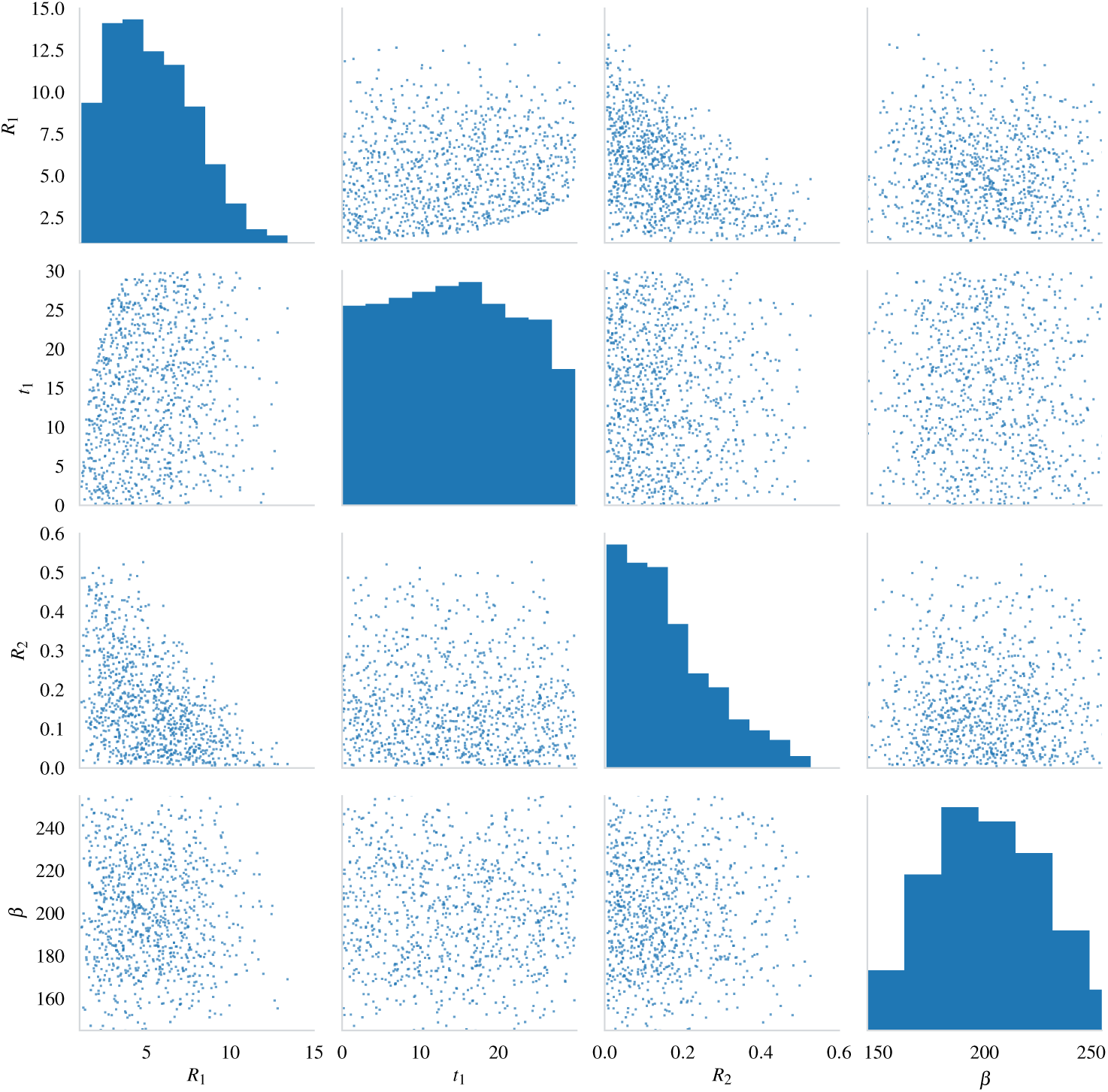
A scatter matrix of samples from the prior.

**Figure S3:**
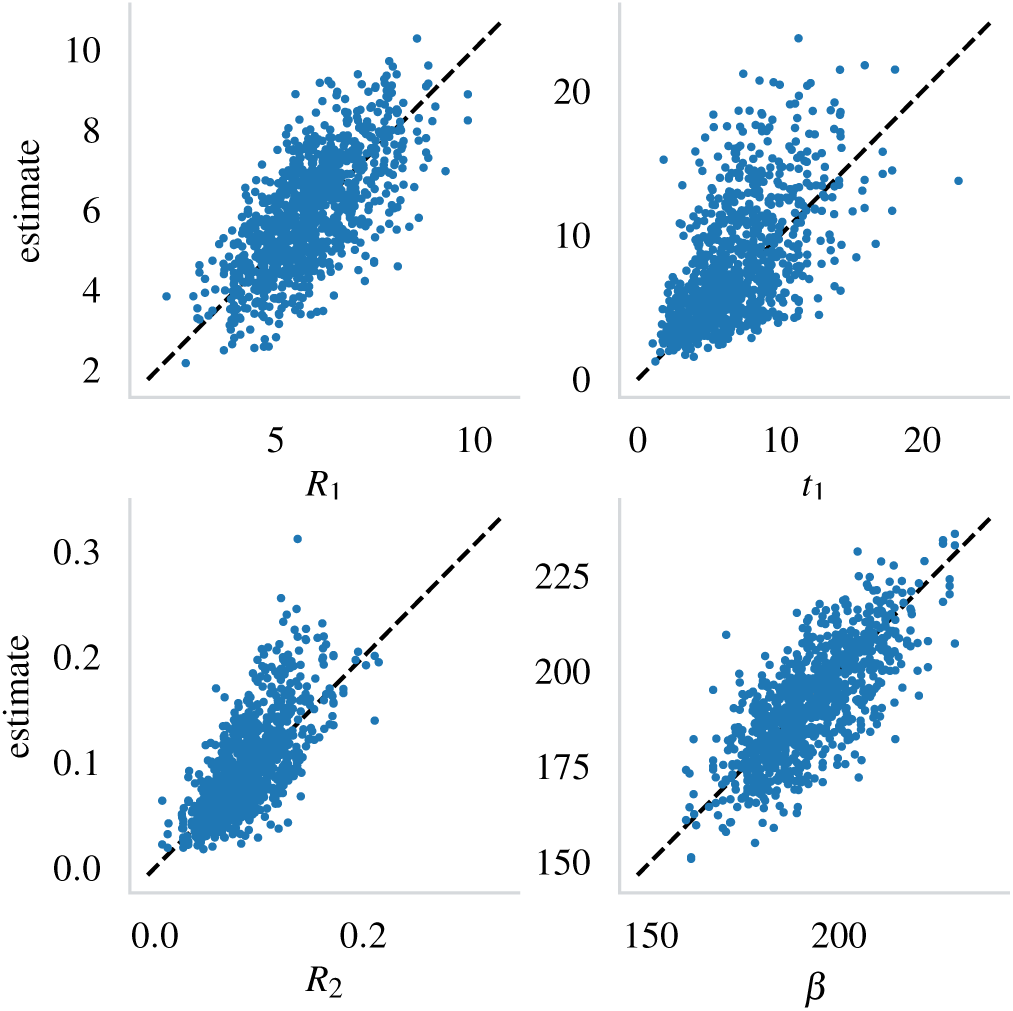
The estimates from the 1000 trials plotted against their true values. The black dashed line shows the 1:1 correspondence.

## Footnotes

1 https://github.com/lintusj1/tb-model

